# Retrotransposon-driven environmental regulation of *FLC* leads to adaptive response to herbicide

**DOI:** 10.1101/2023.09.06.556499

**Authors:** Mathieu Raingeval, Basile Leduque, Pierre Baduel, Alejandro Edera, Fabrice Roux, Vincent Colot, Leandro Quadrana

## Abstract

The mobilization of retrotransposons yields major-effect mutations. Here, we report an adaptive retrotransposon insertion within the first intron of the Arabidopsis floral-repressor locus *FLOWERING LOCUS C (FLC)*. The insertion-mutation augments the environmental sensitivity of *FLC* by affecting the balance between coding and non-coding transcript isoforms in response to environmental threads. We show that this balance is modulated epigenetically by DNA methylation and orchestrated by IBM2, a factor involved in the processing of intronic heterochromatin. The stress-sensitive allele of *FLC* has recently spread across populations subjected to recurrent chemical weeding, and we demonstrate that retrotransposon-driven acceleration of life cycle represents a rapid response to herbicide. Our findings illustrate how retrotransposition can create environmentally-sensitive alleles that facilitate adaptation to anthropogenic disturbances of the environment.

## INTRODUCTION

Environmental changes due to direct human activities have impacted biodiversity at unprecedented pace, putting a countless number of natural populations at evolutionary risk. Cropland expansion, intensified chemical input, and urbanization of natural habitats have favored the replacement of native populations by colonizing ruderal species. Identifying the type, mode of action, and history of mutations enabling adaptation to ruderal habitats is crucial to forecast the impact of ongoing and future anthropogenic activity on biodiversity.

Among the different types of mutations, those that occur spontaneously following DNA replication errors or damage are arguably the best characterized (Mackay, Stone, and Ayroles 2009; Weigel and Nordborg 2015). However, structural variants, notably those produced by the mobilization of transposable elements (TEs), which are DNA sequences that have the ability to self-propagate within and across genomes, account for the largest fraction of varying base pairs among individuals (Michael et al. 2017; Sudmant et al. 2015; Weischenfeldt et al. 2013). TE mobilization has been proposed to have a greater potential than single nucleotide polymorphisms to generate large effect mutations (Schrader and Schmitz 2019), which may be further accentuated by the exquisite sensitivity of some TEs to environmental signals (Casacuberta and González 2013). Despite these considerations, the actual contribution of new TE insertions to adaptive walks remains controversial.

Previous analyses of TE insertions polymorphisms based on population genomics data for hundreds of accessions of the model species *Arabidopsis thaliana* revealed that the first intron of the *Flowering Locus C* (*FLC*) gene is the most frequent target of TE insertions in nature, with up to 16 independent TE-containing alleles (Quadrana et al. 2016; Baduel et al. 2021). *FLC* encodes a MAD-box transcription factor that negatively regulates floral transition by impairing the expression of flowering signals. Vernalization (i.e. overwintering or long cold exposure at the seedling stage) represses *FLC* and this repression is maintained epigenetically upon the return to warmth, allowing plants to flower in spring (Bastow et al. 2004). However, with few exceptions (Qüesta et al. 2016), we currently have limited knowledge about how mutations at *FLC* modulate flowering time variation in nature.

## RESULTS

### A recent retrotransposon insertion within FLC modulates flowering response

Using long-read sequencing data, we assembled a highly contiguous, telomere-to-telomere, genome assembly of the Arabidopsis accession Ag-0 (Supplementary Figure 1), which carries a retrotransposon insertion within *FLC (Quadrana et al. 2019)*. Our assembly confirmed the localization and identity of a full-length *ATCOPIA78*/*ONSEN* retrotransposon insertion within *FLC*’s first intron (Figure 1a). Based on the nucleotide divergence between retrotransposon’s LTRs (SanMiguel et al. 1998) we estimated that this insertion is less than 0.16 million years old (MYO). The Ag-0 genome has two additional full-length *ONSEN* insertions, located in chromosomes 1 and 3. LTRs of these copies were significantly divergent (2.27% and 0.66%, respectively), suggesting they inserted ∼2.27 and ∼0.66 MYO, respectively. Consistently, these insertions are respectively fixed or at intermediate frequency (14%) among 1069 natural Arabidopsis accessions previously characterized (Baduel et al. 2021). The insertion within *FLC* is more closely related to *ONSEN* copies carried by the reference accession Col-0 than the ones on chromosome 1 and 3 of Ag-0 (Figure 1b), suggesting that none of these two served as donors for the insertion within *FLC*. To characterize the origin of that insertion, we set out to identify accessions carrying the same *FLC* haplotype as Ag-0 using SNP data available for the 1069 natural accessions (1001 Genomes Consortium 2016). Twenty-seven accessions, mostly from northern latitudes, carry the Ag-0 haplotype (Hap1) but lack the retrotransposon insertion (Figure 1c). Based on the number of SNPs accumulated in the haplotype block we estimated that the transposon insertion within *FLC* occurred less than 246 years ago. Considering that Ag-0 was collected by Denis Ratcliffe almost 60 years ago from a railway ballast (Kranz and Kirchheim 1987), it is most likely that the insertion within *FLC* took place during the XIX century.

**Figure 1.**
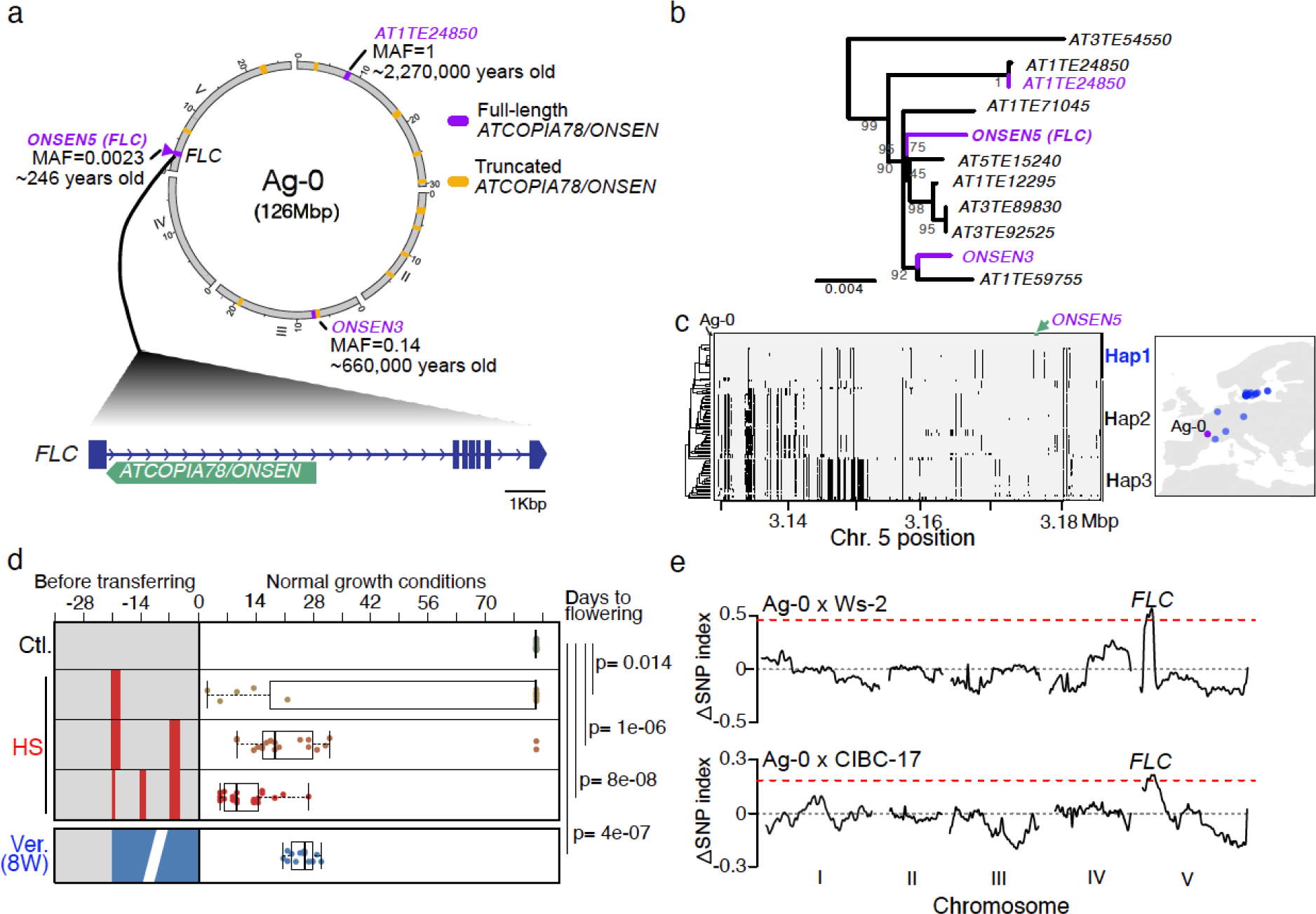
A recent retrotransposon insertion within *FLC* modulates flowering response. **a.** Circos plot of Ag-0 genome assembly with full-length (violet) and truncated (yellow) ATCOPIA78/ONSEN insertions. The localization, minor allele frequency (MARF) across 1069 natural accessions, and estimated age of full-length insertions are indicated. **b**. Phylogenetic tree of full-length *ONSEN* insertions in Ag-0 and the reference genome Col-0. Bootstrap probabilities are shown beside the branches. **c**. Polymorphisms along the *FLC* haplotype block between accessions with related haplotypes. Ag-0 accession and location of the *ONSEN* insertion are indicated. Geographic distribution of accessions sharing the Ag-0 haplotype of *FLC* (Hap1) is shown. **d**. Boxplots of days to flowering of Ag-0 plants grown under control conditions (Ctl.), subjected to eight weeks of vernalization (8W Vern.), or subjected to a single, or repeated heat shock (HS) at the seedling stage. Timing of the HSs are indicated by red bars. Statistical significance for differences were obtained using a two-sided MWU test. **e**. Genome-wide Δ(SNP index) plot of F2 (Ag-0 x Ws-2 and Ag-0 x CIBC-17) bulk segregant analysis of flowering responses to heat-shock. Red line indicates the genome-wide 1% threshold. Localization of *FLC* locus is indicated.

Numerous TE insertions within *FLC* have been identified in Brassicaceae species, most of which produce loss-of-function alleles (Quadrana 2020). To test if the TE-containing *FLC* allele in Ag-0, which has one of the longest (8435nt-long) introns described in Arabidopsis (Supplementary Figure 2), is functional and subjected to its canonical regulation, we grew Ag-0 plants under standard long-days conditions or eight weeks of vernalization. Vernalized plants flowered soon after the treatment (28±1.39 days after vernalization), while control plants flowered more than 120 days after germination (Figure 1d). Consistently, *FLC* expression was initially high and strongly reduced following vernalization (Supplementary Figure 3). Vernalization-induced down-regulation of *FLC* was maintained at least four weeks after returning plants to warm temperatures, similarly to the Northern Sweden ecotype Lov-1 (Coustham et al. 2012) that requires long cold exposure to saturate the vernalization requirement (Supplementary Figure 3). These results confirm that the *FLC* allele carried by Ag-0 is functional and characterized by a canonical vernalization response.

Because *ONSEN* retrotransposons are induced by heat shock (HS) (Tittel-Elmer et al. 2010), we tested the flowering response of Ag-0 to single or multiple HS (Figure 1d). Compared to the late flowering response observed when plants are grown under control conditions, plants subjected to heat stresses flowered much earlier (40-70 days after germination). This acceleration in flowering correlates with the severity and number of heat-shocks. For instance, while only 30% of plants subjected to a single day of heat shock flower less than 50 days after germination, all plants subjected to three heat shocks flowered ∼40 days after germination.

To determine the genetic architecture of the flowering response to HS, we generated an F2 population by crossing Ag-0 with two accessions with different degree of vernalization response (Ws-2 and CIBC17). Bulk segregant analysis of F2 plants flowering early in response to HS identified a single peak of association spanning the Ag-0 allele of *FLC* in both crosses (Figure 1e), further suggesting that the TE-containing allele of this gene is causal for the HS-induced flowering response.

### Heat-induced retrotransposon activation impairs FLC activity

To investigate the molecular mechanisms underpinning the flowering response of *FLC* to HS, we performed RNA-seq experiments on Ag-0 seedlings grown under control or HS conditions. *FLC* was highly expressed in control plants (Figure 2a), in agreement with the strong vernalization requirement observed for this accession. *ONSEN* was strongly induced in response to HS (Figure 2a), consistent with previous work reporting activation of this TE in several Arabidopsis accessions (Masuda et al. 2016). Alongside *ONSEN* reactivation, the first intron of *FLC* is significantly retained and overall *FLC* expression was markedly reduced in response to stress. RT-qPCR confirmed these observations (Figure 2b) (Quadrana et al. 2019). We did not find, however, significant readthrough transcription from the *ONSEN* insertion towards the first exon of *FLC,* suggesting that nonsense-mediated degradation, rather than retrotransposon-derived antisense transcripts likely cause the observed downregulation of *FLC*. Moreover, the canonical *FLC* antisense transcript *COOLAIR*, which mediates the epigenetic silencing of *FLC* (Liu et al. 2010), is also downregulated in response to heat stress (Supplementary Figure 4), excluding its participation in the reduction of *FLC* expression. In addition, expression of *FLM*, another negative regulator of flowering time known to be downregulated in response to warm temperatures (Posé et al. 2013), is not altered in response to HS (Supplementary Figure 4). Crucially, HS-induced downregulation of *FLC* was accompanied by a strong induction of direct targets of *FLC (Deng et al. 2011)*, such as the major flowering integrators *FLOWERING LOCUS T (FT)*, *SUPPRESSOR OF OVEREXPRESSION OF CONSTANS 1 (SOC1)*, and *AGAMOUS LIKE 24 (AGL24)* (Figure 2b and Supplementary Figure 4). In combination, these results indicate that environmental reactivation of *ONSEN* impairs *FLC* expression, inducing the expression of flowering integrators that trigger flowering transition.

**Figure 2.**
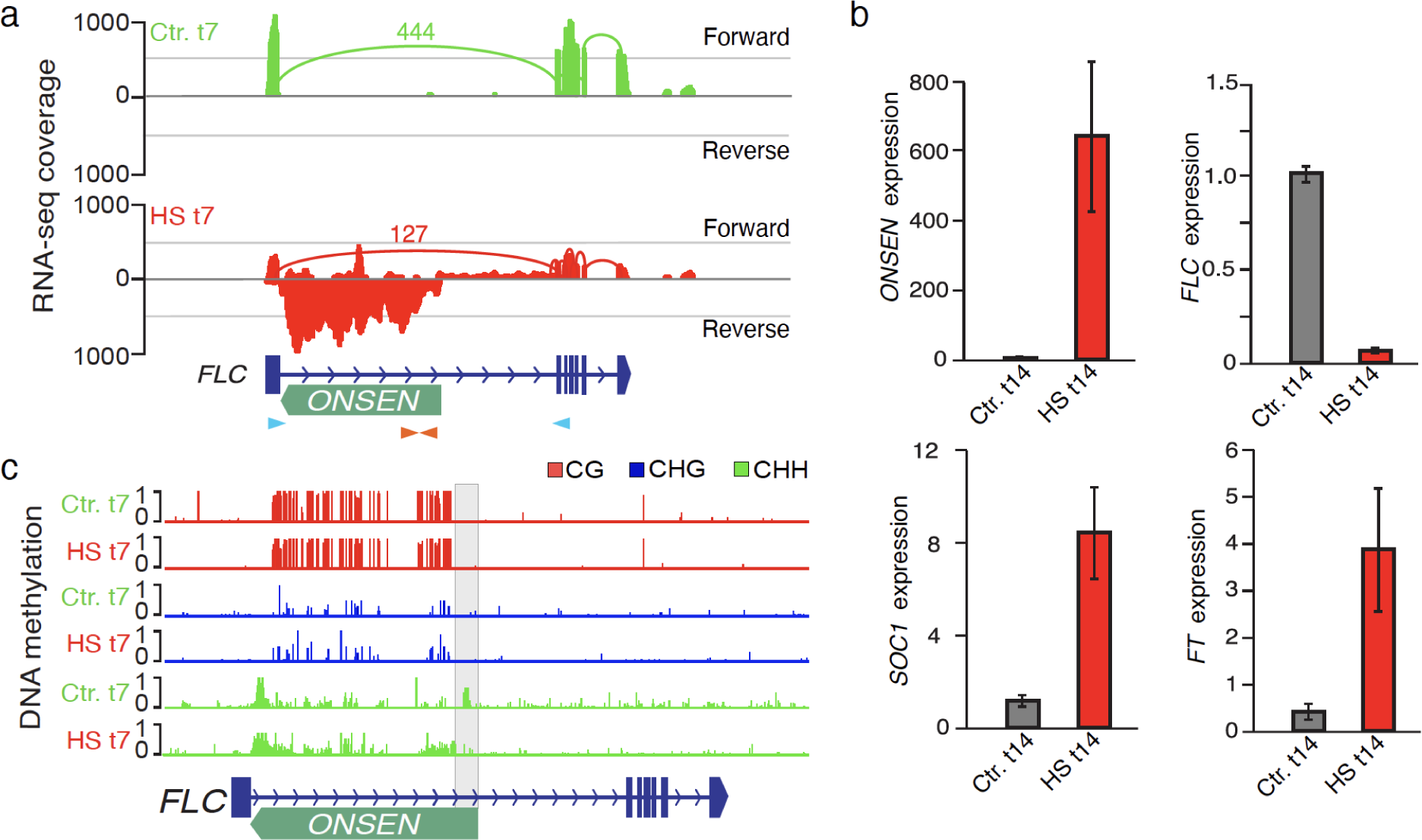
Heat-induced retrotransposon activation impairs *FLC* activity. **a**. Genome browser view of normalized RNA-seq coverage at *FLC* locus in Ag-0 plants seven days after a heat-shock (HS t7) or control treatment (Ctl. t7). Light blue and orange arrowheads represent the location of primers used to quantify *FLC* and *ONSEN* expression, respectively. **b**. Relative expression levels of *ONSEN*, *FLC*, *SOC1*, and *FT* in Ag-0 plants 14 days after a heat-shock (HS t14) or control treatment (Ctl. t14). Data are mean ± s.d. (n>10 samples, two biological experiments) and statistical significance for differences was obtained using the MWU test. **c**. Genome browser tracks showing local CHH hypomethylation of the 5’ end of the *ONSEN* insertion (gray box) in response to heat-shock (HS t7).

Experimental loss of heterochromatin modifications, including DNA methylation, can affect the expression of TE-containing genes (Saze et al. 2013; Berthelier et al. 2023). However, whether HS-induced reactivation of *ONSEN* is associated with its hypomethylation is still unclear (Tittel-Elmer et al. 2010; Sun et al. 2020), likely due to the difficulty to investigate DNA methylation of such repetitive sequences. In contrast to the reference Arabidopsis genome Col-0, which contains at least four nearly identical full-length *ONSEN* copies, Ag-0 has only three, and significantly divergent, copies (Figure 1b). Indeed, bisulfite-sequencing (Bi-seq) data obtained from control Ag-0 plants provide robust DNA methylation information across the complete *ONSEN* annotation and revealed extensive non-CG methylation of the LTR sequences (Figure 2c). In response to HS, the 5’ LTR of *ONSEN,* which contains heat-shock response elements and acts as a promoter of transcription (Pietzenuk et al. 2016), becomes strongly and rapidly hypomethylated, indicating that transcriptional reactivation of *ONSEN* is accompanied by DNA methylation loss.

### DNA methylation-dependent modulation of FLC

To test directly the role of DNA methylation in the modulation of flowering response in Ag-0, we generated CRISPR-Cas9-mediated mutants of the SWI2/SNF2 chromatin remodeler DDM1, which is required to maintain DNA methylation in Arabidopsis (Jeddeloh, Stokes, and Richards 1999). After two generations with the CRISPR-Cas9 cassette, transgene-free offspring were propagated and genotyped to identify plants carrying mutations between the end of exon 6 and the beginning of exon 7 of *DDM1*, where guide RNAs were targeted (Supplementary Figure 5a). Two lines (*ddm1-5* and *ddm1-6*) were obtained in this way (Figure 3a). Target sequencing revealed that both mutants carry a similar frameshift mutation in exon 6 that generates multiple premature stops within the ATPase domain of DDM1 (Supplementary Figure 5b). DNA methylation levels of *ONSEN* were strongly reduced in the CRISPR-mutant compared to wild-type Ag-0 plants (Figure 3b), confirming the loss of DDM1 activity in these plants. Strikingly, compared to the late flowering phenotype of control Ag-0 plants, *ddm1* plants flower 25±6 days after germination (Figure 3a). We then analyzed the expression of canonical as well as intron-retention (IR) transcript isoforms of *FLC. ddm1*-induced hypomethylation of *ONSEN* was accompanied by a dramatic downregulation of *FLC* expression and production of IR-*FLC* transcript isoforms (Figure 3c and Supplementary Figure 5b). Notably, *ONSEN* expression was not altered in *ddm1* compared to wild-type Ag-0 plants (Supplementary Figure 5c), In combination, these results established that it is the hypomethylation of *ONSEN,* rather than its transcriptional activation, that modulates *FLC* expression and floral transition in Ag-0 (Figure 3d).

**Figure 3.**
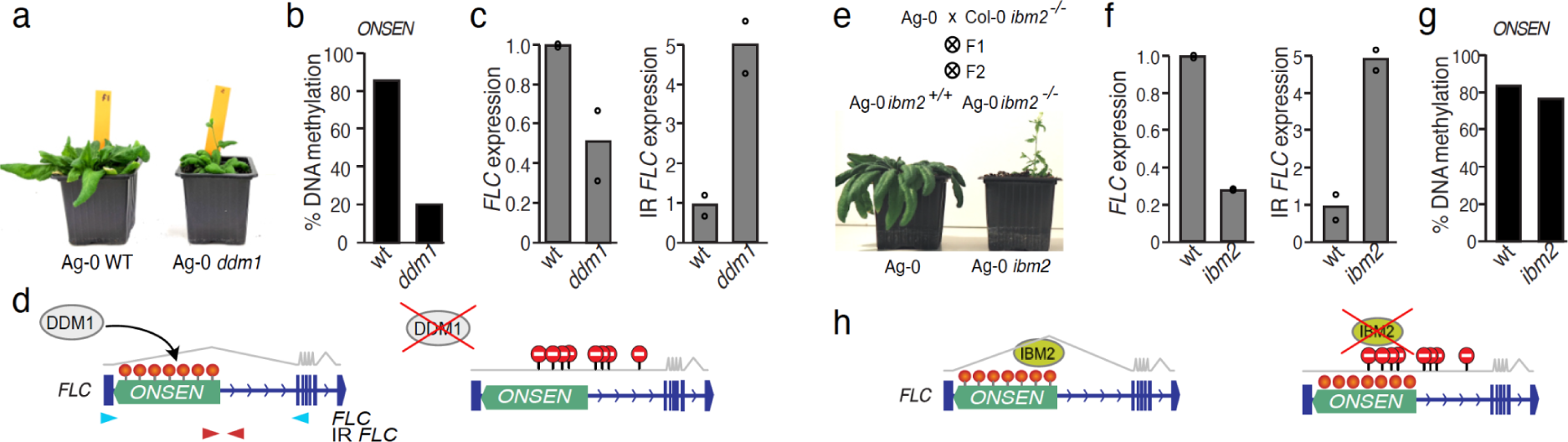
DNA methylation-dependent modulation of *FLC*. **a.** Wild-type (WT) and *ddm1* mutant Ag-0 plants. **b.** DNA methylation levels of *ONSEN in wild-type and ddm1* mutant Ag-0 plants**. c.** Relative expression levels of full-length and intron-retention (IR) transcript isoforms of *FLC*. Primers’ locations are indicated in panel d. Data are mean ± s.d. (n>10 samples, two biological experiments) and statistical significance for differences was obtained using the MWU test. **d.** Model summarizing the function of DDM1 in the modulation of DNA methylation (red lollipops) and appropriate processing of *FLC* transcripts. **e.** Wild-type (WT) and *ibm2* mutant Ag-0 plants. **f.** Relative expression levels of full-length and intron-retention (IR) transcript isoforms of *FLC.* Primers’ locations are indicated in panel d. Data are mean ± s.d. (n>10 samples, two biological experiments) and statistical significance for differences was obtained using the MWU test **g.** DNA methylation levels of *ONSEN in wild-type and ibm2* mutant Ag-0 plants. **h.** Model summarizing the function of IBM2 in the appropriate processing of *FLC* transcripts.

Processing of gene transcripts produced from intronic heterochromatin is mediated by a protein complex comprising the RNA binding protein *INCREASE IN BONSAI METHYLATION 2* (*IBM2*) (Saze et al. 2013). To test the involvement of IBM2 in the expression of *FLC* in Ag-0, we introduced the *ibm2-4* mutant allele (Deremetz et al. 2019) into Ag-0 plants. We selected F3 plants carrying the Ag-0 alleles of *FLC* in combination with either the WT or the mutant *IBM2* alleles (Figure 3e). These plants (Ag-0 *ibm2^-/-^* and Ag-0 *ibm2^+/+^,* respectively) were grown under control conditions to measure flowering time. Ag-0 *ibm2^-/-^* mutant plants flowered 22 +/-10 days after germination, while all Ag-0 *ibm2^+/+^* control plants flowered more than 120 days after germination (Figure 3e). Expression analysis showed that acceleration of flowering in Ag-0 *ibm2^-/-^* plants was accompanied by a significant reduction in *FLC* expression and induction of IR-*FLC* transcripts isoforms (Figure 3f). As expected, misregulation of *FLC* in *ibm2* plants leads to higher *FT* expression levels (Supplementary Figure 6), further supporting that failed processing of the *ONSEN*-containing intron of *FLC* transcripts, such as in response to HS or *ddm1*-induced hypomethylation, underlie the early flowering of Ag-0. Importantly, DNA methylation of the *ONSEN* insertion was not affected in Ag-0 *ibm2^-/-^* plants (Figure 3g), indicating that IBM2 functions downstream of DNA methylation. Altogether, these results demonstrate that the environmental modulation of the TE-containing allele of *FLC* is mediated by IBM2’s methylation-dependent transcript processing (Figure 3h).

### Retrotransposon-mediated acceleration of life cycle in response to herbicide

Given the key adaptive role of *FLC* in aligning flowering time with seasons, the retrotransposon-driven HS impairment of *FLC* expression may reflect a more general ability of Ag-0 plants to flower early in response to specific environmental conditions (Quadrana et al. 2019). To investigate whether the TE-containing allele of *FLC* has spread across populations experiencing similar climates as that of Ag-0, we collected in spring 2019 and 2021, 74 Arabidopsis accessions across a 160km2 area near the city of Argentat in central France (Figure 4a and table S1), where Ag-0 was initially collected in the 50s (Kranz and Kirchheim 1987). The newly collected accessions were growing in diverse habitats, including brownfields, sidewalks, railways, and field margins. Genotyping showed that 23% of the accessions carry the retrotransposon insertion in *FLC*, demonstrating that this allele has been maintained in the area for the past 60 years. Intriguingly, none of the populations from Argentat, or nearby cities, carry the insertion (Figure 4a), as the railway where Denis Ratcliffe initially collected Ag-0 was dismantled in 1970 and the zone became heavily urbanized since then. Conversely, the insertion is frequent (∼40%) in Arabidopsis populations nearby the city of Archignac and Turenne, yet carrier and non-carrier populations share the same climate niche (Figure 4b). Therefore, the retrotransposon-containing *FLC* allele more likely plays a role in adaptation to local environments, rather than providing an adaptive response to warmer climates as previously proposed (Quadrana et al. 2019). Consistent with this possibility, carriers of the TE-containing allele of *FLC* were found most often along railways and field margin strips (Figure 4c), two environments exposed to intense chemical weeding (Marshall and Moonen 2002; Antuniassi, Velini, and Nogueira 2004). Selection of weeds in such environments has been linked to mutations in several flowering pathways, notably *FLC (Baduel et al. 2018)*, enabling plants to cycle faster and complete a full life-cycle in high-stress and high-disturbance environments (Kreiner et al. 2022). To test this hypothesis, we investigated the presence of the retrotransposon-containing allele in 165 natural populations of Arabidopsis collected across south-west of France in 2015 (Frachon et al. 2018). Remarkably, and despite its recent origin, the retrotransposon-containing allele of *FLC* is present in 56% of these populations (Figure 4a, table S2). Carrier populations were significantly enriched in railways, and, to a lesser extent, field margins, backyards, and silo sites (Figure 4d), supporting a role for this allele in the response to chemical weeding. To determine if this allele is under positive selection, we quantified haplotype-lengths using the Extended Haplotype Homozygosity (EHH) test (Sabeti et al. 2002), which detects recent selective sweeps associated with *de novo* mutations. This analysis showed that haplotypes carrying retrotransposon insertions are more frequent and much longer than ancestral haplotypes (Figure 4e), indicating recent or ongoing positive selection of the insertion. In combination, these results demonstrate that the *de novo* retrotransposon insertion within *FLC* is likely involved in adaptation to herbicide-heavy environments.

**Figure 4.**
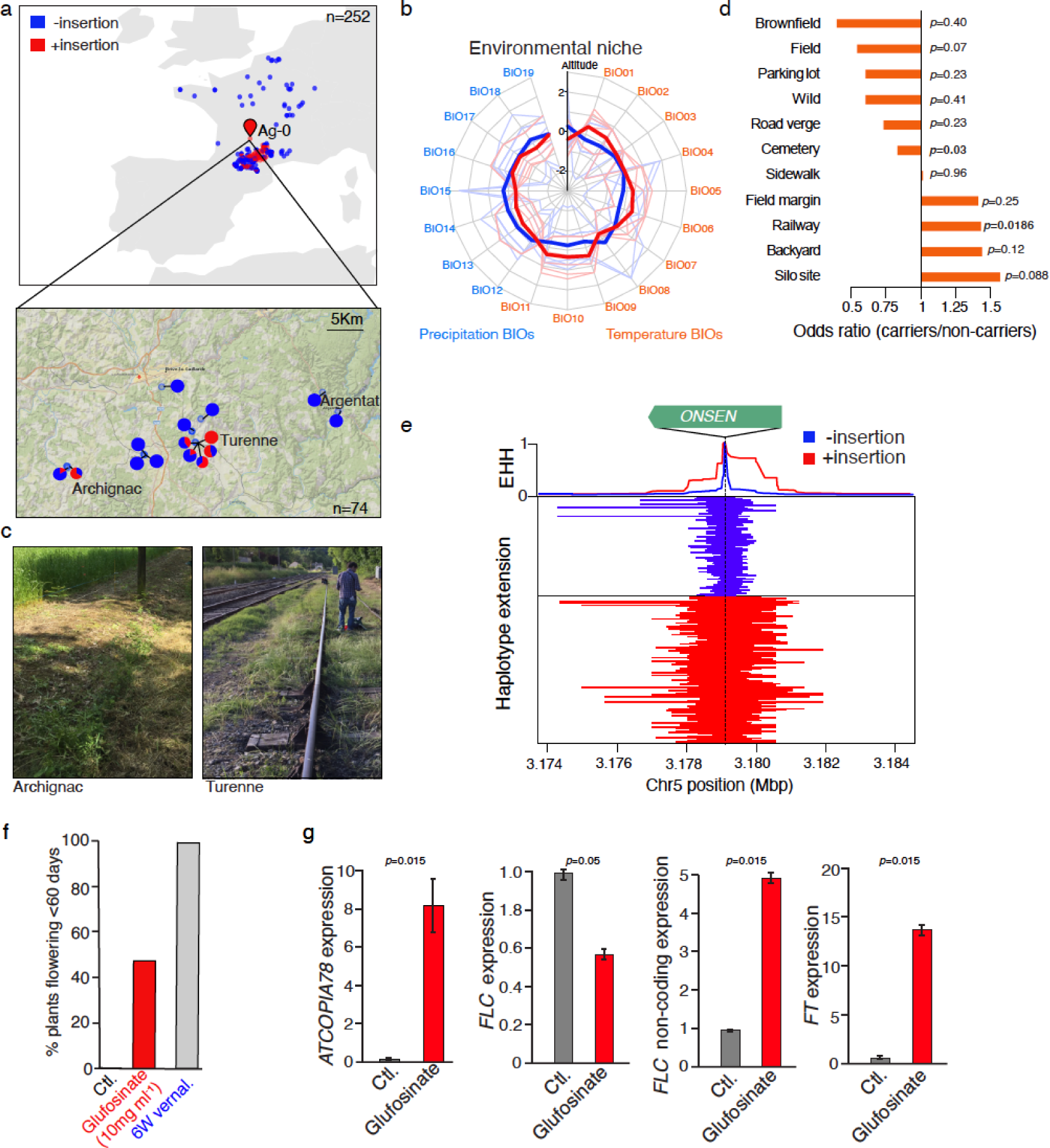
Retrotransposon-mediated acceleration of life cycle in response to herbicide. **a.** Geographic distribution of Arabidopsis populations collected across France. A detailed view of populations collected nearby the Argentat is shown. Carriers and non-carriers of the *ONSEN*-containing allele of *FLC* are indicated in red and blue, respectively. **b**. Climate niche description of populations nearby Argentat. Carrier and non-carrier populations are indicated in red and blue, respectively. **c**. Picture of collection sites where carrier populations were found. **d**. Relative enrichment (odds ratio) of carrier populations from different local environments. Statistical significance for differences was obtained using the chi-square test. **e**. Extended haplotype homozygosity (EHH) at varying distances from the *ONSEN* insertion site for carriers and non-carriers. **f**. Percentage of plants flowering within 60 days after germination grown under control conditions, subjected to sub-lethal doses of Glufosinate, or vernalized. Statistical significance for differences was obtained using the chi-square test. **g**. Relative expression levels of *ONSEN*, full-length and intron-retention (IR) transcript isoforms of *FLC*, and FT. Data are mean ±s.d. (n>10 samples, one biological experiment) and statistical significance for differences was obtained using the MWU test.

To test directly this possibility, we sprayed Ag-0 plants with sublethal doses of the broad-spectrum herbicide Glufosinate. Strikingly, at least half of the Ag-0 plants treated with herbicide flowered before 60 days, compared to none of the control plants (Figure 4f). RT-qPCR experiments show that under these conditions *ONSEN* becomes strongly upregulated in response to herbicide (Figure 4g), supporting the notion that this retrotransposon is sensitive to severe stresses (Matsunaga et al. 2015), rather than being induced by specific environmental signals such as HS. Together with retrotransposon activation, the balance between full-length and IR *FLC* transcripts drastically shifts, leading to high expression of the non-functional *FLC* isoform, *FT* induction, and early flowering (Figure 4g). In conclusion, the retrotransposon insertion within *FLC* enables plants to cycle faster in response to chemical spraying, providing a rapid adaptive response of Arabidopsis to modern weed control practices (Supplementary Figure 7).

## DISCUSSION

Agricultural intensification and extensive urbanization, aided by the massification of agrochemicals, has transformed natural habitats at unprecedented pace. How populations react to such drastic environmental changes, and whether it relies on standing variation or de novo mutations, is still a matter of debate. We demonstrate that a gain-of-function retrotransposon insertion within the flowering gene *FLC* enables Arabidopsis plants to cycle faster in response to herbicide-exposure, leading to an escape response to chemical weeding. This escape mechanism may apply to a wide array of stresses, setting them apart from conventional herbicide resistance responses, which entail the accumulation of mutations at genes encoding specific herbicide target enzymes or related metabolic pathways (Kreiner et al. 2022; Kersten et al. 2023). Additionally, while target-site herbicide resistances are typically caused by fixed-effect mutations, the effect of the retrotransposon insertion in *FLC* allele is stress-dependent. Such a type of mutation should reduce fitness costs in the absence of stress, and therefore increase its chance to persist in the population as segregating alleles (Merenciano and González 2023; Baduel et al. 2021). Indeed, this insertion occurred before the discovery of herbicides in the late 1800s, and likely maintained in the population. In line with this scenario, the TE-containing allele of *FLC* is found only at intermediate frequencies across recently collected populations (Figure 4a). Thus, environmentally-dependent fitness effects may facilitate the spreading of TE insertions through balancing selection at the regional scale, providing an alternative explanation for the excess of rare TE insertion alleles observed across natural populations at the worldwide scale (Merenciano and González 2023; Baduel et al. 2021, 2019; Niu et al. 2019).

Our findings provide a compelling example of a recent retrotransposon insertion enabling a rapid evolutionary response to anthropogenic activities. Adaptive TE insertions were previously reported (Casacuberta and González 2013). For instance, the industrial melanism of the British peppered moths was caused by a *de novo* TE insertion within the gene cortex involved, through still unclear mechanisms, in pigmentation of Lepidoptera’s wings (Van’t Hof et al. 2016). Similarly, natural pesticide resistance in *Drosophila melanogaster* was found associated with the presence of a TE insertion within the gene CHKov1, likely affecting a target of organophosphate pesticides (Aminetzach, Macpherson, and Petrov 2005). Unlike these adaptive TE insertions mutations, however, the response reported here relies on the ability of the retrotransposon *ONSEN* to respond to environmental threats. The *ONSEN-*containing allele of *FLC* is reminiscent of the retrotransposon *KARMA* within the gene *MANTLED*, the hypomethylation of which leads to splicing alterations and somaclonal variations in oil palm (Ong-Abdullah et al. 2015). Hypomethylation of KARMA occurs in response to repeated *in vitro* culture. Hence, environmentally- and artificially-induced hypomethylation of intronic TEs can be an important source of phenotypic changes in plants, with implications for adaptation and crop production.

*FLC,* and notably its first intron, is a common target of TE insertions in Arabidopsis and other Brassicaceae species (Quadrana 2020; Li et al. 2023), which adds to the remarkable regulatory complexity of this gene. The large number of TE containing alleles of *FLC* likely reflects the major adaptive role of this gene, which we show can be catalyzed by complex mutations such as the insertion of environmentally-sensitive retrotransposon sequences. The insertion-mutation carried by Ag-0 lies close to the H3K27me3 nucleation region, a key sequence for vernalization response that plays a pivotal role in determining *FLC’s* epigenetic state (Yang, Howard, and Dean 2014). If and how such TE insertion modulates the chromatin-mediated vernalization response of *FLC* warrants further investigation. Similarly, the large number of TE insertions within *FLC* offers a unique opportunity to explore how transposition can contribute to novel gene regulation mechanisms. Our findings reveal how retrotransposition can enable rapid response to environmental threads, with implications for our understanding of the genetic basis of past and future adaptations to anthropogenic activities.

## Acknowledgements

We thank members of the Colot and Quadrana group for discussions and critical reading of the manuscript. We thank members of the Colot group, Mercedes Tkach, Hugo Vergne, and Claire Delamare Deboutteville for helping with the collection of Arabidopsis plants nearby Argentat. We also thank Nicolas Bouché for sharing *ibm2-4* mutant seeds and Fabien Nogué for sharing the pDE-Cas9-dsRed plasmid as well as advice on guide RNA design.

## Authors contributions

LQ conceived the project. MR performed most HS and herbicide experiments, RT-qPCRs, and generated Ag-0 *ibm2* mutants, with additional help from LQ. PB and VC generated Ag-0 *ddm1* mutant lines. BL genotyped Arabidopsis accessions nearby Argentat, generated ONT sequencing data, assembled and annotated Ag-0 genome, with additional help from AE. FR contributed to the analysis of south-west French populations. LQ performed BS-seq, RNA-seq, QTL mapping, genomic and population genomic analyses, interpreted the data, and wrote the manuscript. All the authors read and approved the paper.

## Data and material availability

Original Scripts are available on GitHub. WGBS and RNA-seq of Ag-0 plants are available at the European Nucleotide Archive (ENA) under project XXXX.

## Materials and Methods

### Samples and experimental models

The following *A. thaliana* plants were used: wild-type Ag-0, Ws-2, CIBC-17, and Lov-1 obtained from the NASC collection. The Col-0 *ibm2-4* mutant was obtained from Deremetz et al 2019. *Ibm2* Ag-0 plants were obtained by crossing Col-0 *ibm2-4* with Ag-0. F2 plants were genotyped and *ibm2-4*^-/-^, *FLC^ag-0/ag-0^*, and *FRI^ag-0/ag-0^,* as Col-0 has a week *FRI* allele that impairs *FLC* expression, were selected and propagated till F3 for phenotypic and molecular characterization. Collections of seeds from Arabidopsis plants around Argentat were carried out in May 2019, August 2019, and April 2021. In order to break potential seed dormancy, seeds were kept at least four months at 15 degrees previous to germination. CRISPR-CAS9 *ddm1* mutants of Ag-0 were generated by floral-dip transformation using the binary vector pDE-Cas9-dsRed following reported protocol (Morineau et al. 2017). Briefly, two guide RNAs targeting the end of exon 6 and the beginning of exon 7 respectively (GAATTCTGCGGCACTATCCA and GATAACAAACTTCTGCTGAC) were designed using CRISPOR (http://crispor.tefor.net/crispor.cgi) and synthesized by Invitrogen GeneArt (ThermoFisher) in tandem repeat, each preceded by a U6 promoter and separated by a tracrRNA (GTTTAAGAGCTATGCTGGAAACAGCATAGCAAGTTTAAATAAGGCTAGTCC GTTATCAACTTGAAAAAGTGGCACCGAGTCGGTGCTTTTTTT), and flanked by attB sites within a pDONR221 plasmid. The guide RNA construct was then ligated into the pDE-Cas9-dsRed vector by LR Gateway (ThermoFisher) by selection against ccdB. Transgenic T1 plants were identified by seed fluorescence followed by PCR genotyping of the transgenic cassette (primers: sgU6-F CGGTGCTTTTTTTGGATCC and DsRed-proR TGCTTCTTGCTGAGCTCACA), and then propagated in the heterozygous state (characterized by the production of both fluorescent and non-fluorescent seeds) for a second generation. Non-fluorescent offspring (G1) plants of the T2s were genotyped by PCR (primers ddm1-ex6F CACGCCTTCCATCAATGCAA and ddm1-in7R AGCTGCCACCAGTGTTAACA) and candidate mutants validated by sanger sequencing. Two *ddm1* mutant lines were identified in this way. Heat shock stresses were performed as previously described (Quadrana et al. 2019). Herbicide treatments were performed on two weeks old plants by spraying 10mg/ml Glufosinate once per week, during three weeks. Unless stated otherwise, all plants were grown in long-days (16h:8h light:dark) at 23°C. Vernalization treatments were performed by growing ten days old plants in short days at 6C for eight weeks.

### De novo assembly and annotation of Ag-0

High-molecular weight genomic DNA of two weeks old Ag-0 plants was extracted as previously reported (Mayjonade et al. 2016) with minor modifications. Briefly, plant material was grinded under liquid nitrogen and 100mg of powder was incubated in 1 ml of lysis buffer (PVP40 1%, NaCl 500mM, Tris ph8 100mM, EDTA 50mM, SDS1.25%, Sodium bisulfite 1%, DRR5mM) during 30 minutes at 50 degrees. DNA was cleaned using KOAc precipitations (1.3M, pH7.5) and DNA purified using 0.05X of seraMAG beads. ONT library (LSK1010) was prepared using 1 ug of HMW DNA and loaded in a R9.4 Minion flow-cell. Sequencing was performed in a Mk1C device during 72hs. Basecalling was performed using Guppy (V) with the HAC model. Long-read fastq files were used for *de novo* genome assembly using CANU (Koren et al. 2017). low error rate Illumina short reads from Ag-0 (1001 genomes project) were used for sequence polishing using pilon software (Walker et al. 2014). Chromosome scaffolding was achieved by running RagTag (Walker et al. 2014; Alonge et al. 2019) with default parameters. TEs and repetitive sequences were annotated using REPET (Flutre et al. 2011).

### Mapping by sequencing of HS-induced flowering

Ag-0 plants were crossed with Ws-2 or CIBC-17, which show different degrees of vernalization response. Derived F1 plants were backcrossed. Approximately 100 F2 plants were germinated in solid medium (½ MS without sugar) and subjected to a 48h HS one week after germination. Plants were transferred to soil after one week of recovery. Flowering plants were sampled 56 days after germination (n=6 and n=20 for Ag-0 x Ws-2 and Ag-0 x CIBC17, respectively) and polled for DNA extraction by CTAB. DNA was also extracted from pooled leaves of F2 plants as control. One μg of total DNA was used to construct libraries and sequencing (paired-end 100nt reads) by BGI Tech Solutions (Hong Kong). Approximately 20M pair-end reads were obtained for each sample and aligned to the Ag-0 genome assembly using Bowtie2 with default parameters. Reads mapping to multiple locations or duplicates were removed using samtools with parameter-q 5 and Picard MarkDuplicates with default parameters (REMOVE_ DUPLICATES = true), respectively. SNPs were called using bcftools mpileup and bcftools call commands. Variant calls with quality lower than 30 or covered by less than 10 reads were removed. Allele frequency of non-reference alleles were calculated and averaged through windows of 200Kbp, with 100Kbp steps. The difference between allele frequencies of HS-induced flowering F2 plants and control F2 plants were calculated and plotted genome wide. Windows with the highest 1% Δ(SNP index) were considered as significant.

### Transcriptomic analysis

RNA from Ag-0 plants was isolated using Rneasy Plant Minikit (Qiagen) according to the supplier’s instructions. One μg of total RNA was used to construct directional libraries and sequencing (paired-end 100nt reads) by BGI Tech Solutions (Hong Kong). About 20M 100nt-long pair-end reads were obtained per sample. Expression level was calculated by mapping reads using STAR v2.5.3a63 on the Ag-0 reference genome with the following arguments--outFilterMultimapNmax 50--outFilterMatchNmin 30--alignSJoverhangMin 3--alignIntronMax 10000. Duplicated pairs were removed using picard MarkDuplicates.

### Quantification of expression by RT-qPCR

RNA was extracted using the RNeasy plant mini kit (Qiagen) from plants. Contaminating DNA was removed using Turbo DNase (Invitrogen) following manufacturer instructions. 1ug of RNA was used for RT using SuperScriptIV (Invitrogen). RT-qPCR was performed using LightCycler 480 SYBR Green I Master mix (Roche) and run in a LightCycler 480 System (Roche). Primer sequences are provided in table S3. RT-qPCR expression levels relative to the *PP2A* housekeeping gene were calculated using the 2^−ΔΔCt^ method. DNA methylation of *ONSEN* was performed by McrBC-digestion followed by qPCR as described before and using the primers listed in table S3.

### Methylome analysis

DNA from Ag-0 plants was extracted using a standard CTAB protocol. Bisulfite conversion, BS-seq libraries and sequencing (paired-end 100nt reads) were performed by BGI Tech Solutions (Hong Kong). Bioinformatic analysis was performed as previously described (Sasaki et al. 2022). Paired-end reads were trimmed using Trimmomatic program (version 0.33) with following parameters “ILLUMINACLIP:TruSeq3-PE.fa:2:30:10 LEADING:3 TRAILING:3 SLIDINGWINDOW:4:15 MINLEN:36” (Bolger, Lohse, and Usadel 2014). Mapping of trimmed sequences to Ag-0 assembled genome, removal of identical reads, and counting of methylated and unmethylated cytosines were performed by Bismark ver. 0.15.0 (Krueger and Andrews 2011).

### Natural Arabidopsis populations from south-west of France

To call presence/absence of the *ONSEN* insertion within FLC, Illumina re-sequencing data from (Frachon et al. 2018) was aligned to 150bp sequence spanning each insertion extremity of the *ONSEN* insertion within *FLC* from the Ag-0 assembly, as well as to the empty insertion site of the reference Col-o genome (TAIR10). Reads mapping uniquely (qual greater than 24) and entirely (CIGAR string lacking any S) to each extremity and the empty insertion site were counted and compared to estimate the TE insertion allele frequency. Local environments at each collection site were retrieved by inspecting satellite images and aerial pictures available at GEOportail (https://www.geoportail.gouv.fr/carte). This process was performed without knowing the presence/absence information. To investigate the presence of selective sweeps, the same re-sequencing data (Frachon et al. 2018) was aligned to the Arabidopsis reference genome TAIR10 using Bowtie2 with default parameters. Reads mapping to multiple genomic locations or duplicates were removed using samtools with parameter-q 5 and Picard MarkDuplicates with default parameters (REMOVE_ DUPLICATES = true), respectively. SNPs were called using bcftools mpileup and bcftools call commands. Variant calls with quality lower than 30 or covered by less than 10 reads were removed. Biallelic SNPs around the *FLC* locus were extracted using bcftools view-v snps-M 2 and phased using beagle (v5.2). Extended haplotype homozygosity (EHH) around the *ONSEN* insertion within *FLC* was calculated using selscan (Szpiech and Hernandez 2014) with the –ehh option.

