## Supplementary material for "Retrotransposon-driven environmental regulation of *FLC* leads to adaptive response to herbicide"

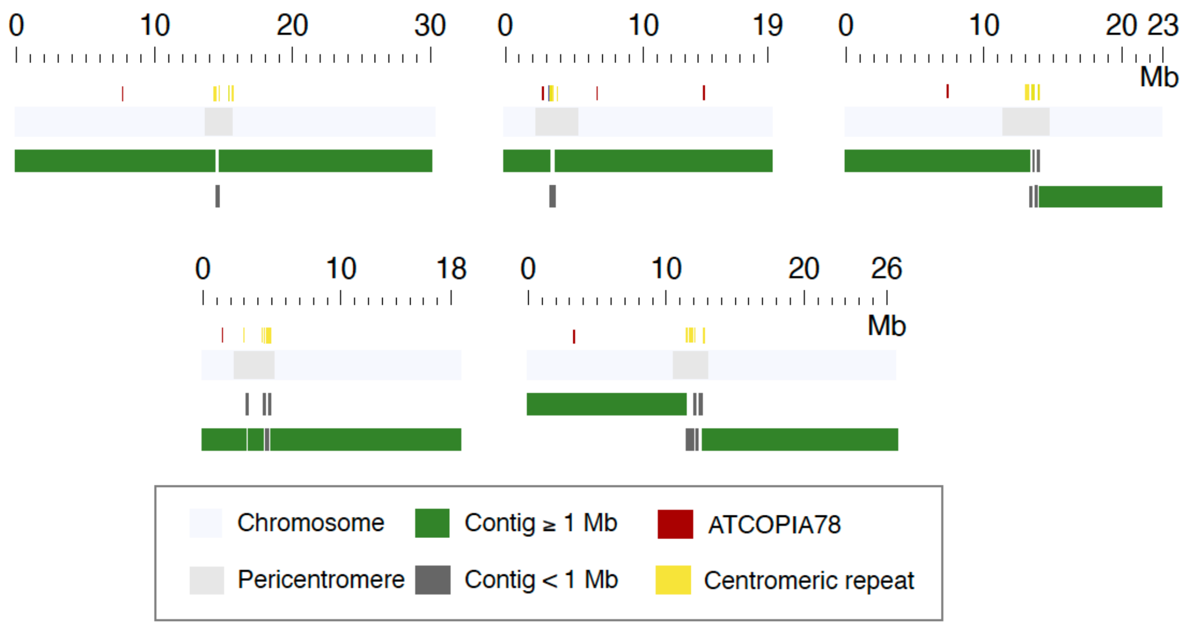


**Supplementary Figure 1. Chromosome-level genome assemblies of Ag-0 accession.** The light gray bars outline each of the chromosomes, whereas the dark gray inlays show the extent of each of the pericentromeric regions. The contig arrangements of the chromosome assemblies is shown in green for contigs > 1 Mb and dark grey for contigs < 1 Mb. The location of centromeric tandem repeat arrays and *ATCOPIA78* insertions within the assemblies are marked by yellow and red boxes above each of the chromosomes.

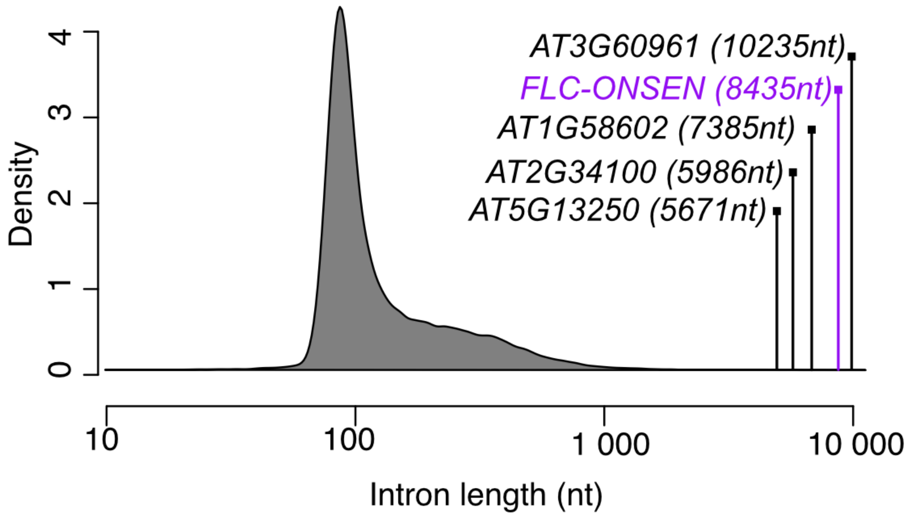


**Supplementary Figure 2. Intron length distribution of all genes annotated in TAIR10**. The length of retrotransposon-containing intron of FLC in Ag-0 is indicated in purple.

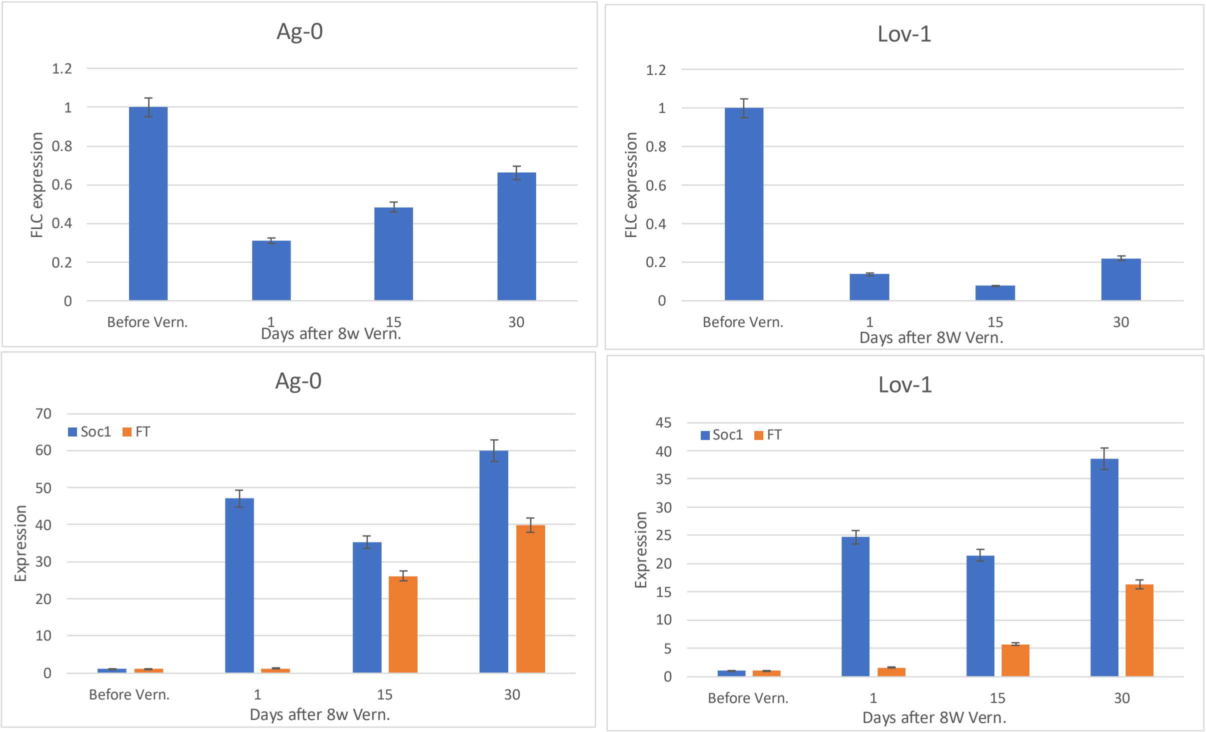


**Supplementary Figure 3. Expression levels of key flowering genes in response to vernalization for Ag-0 and Lov-1 accessions**. Relative expression levels of *FLC*, SOC1, and FT in Ag-0 and Lov-1 plants before and after one, 15, or 30 days of eight-weeks vernalization treatment.

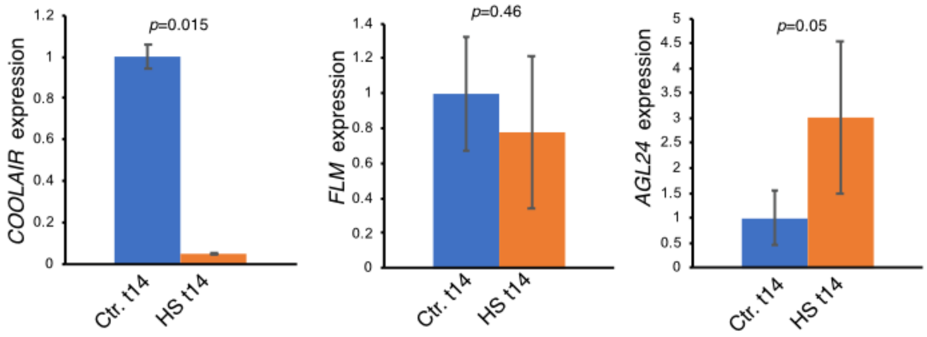


**Supplementary Figure 4. Expression levels of key flowering genes in response to HS for Ag-0**. Relative expression levels of *COOLAIR*, *FLM*, and *AGL24* in Ag-0 plants 14 days after a heat-shock (HS t14) or control treatment (Ctl. t14). Data are mean ± s.d. (n>10 samples, two biological experiments) and statistical significance for differences was obtained using the MWU test

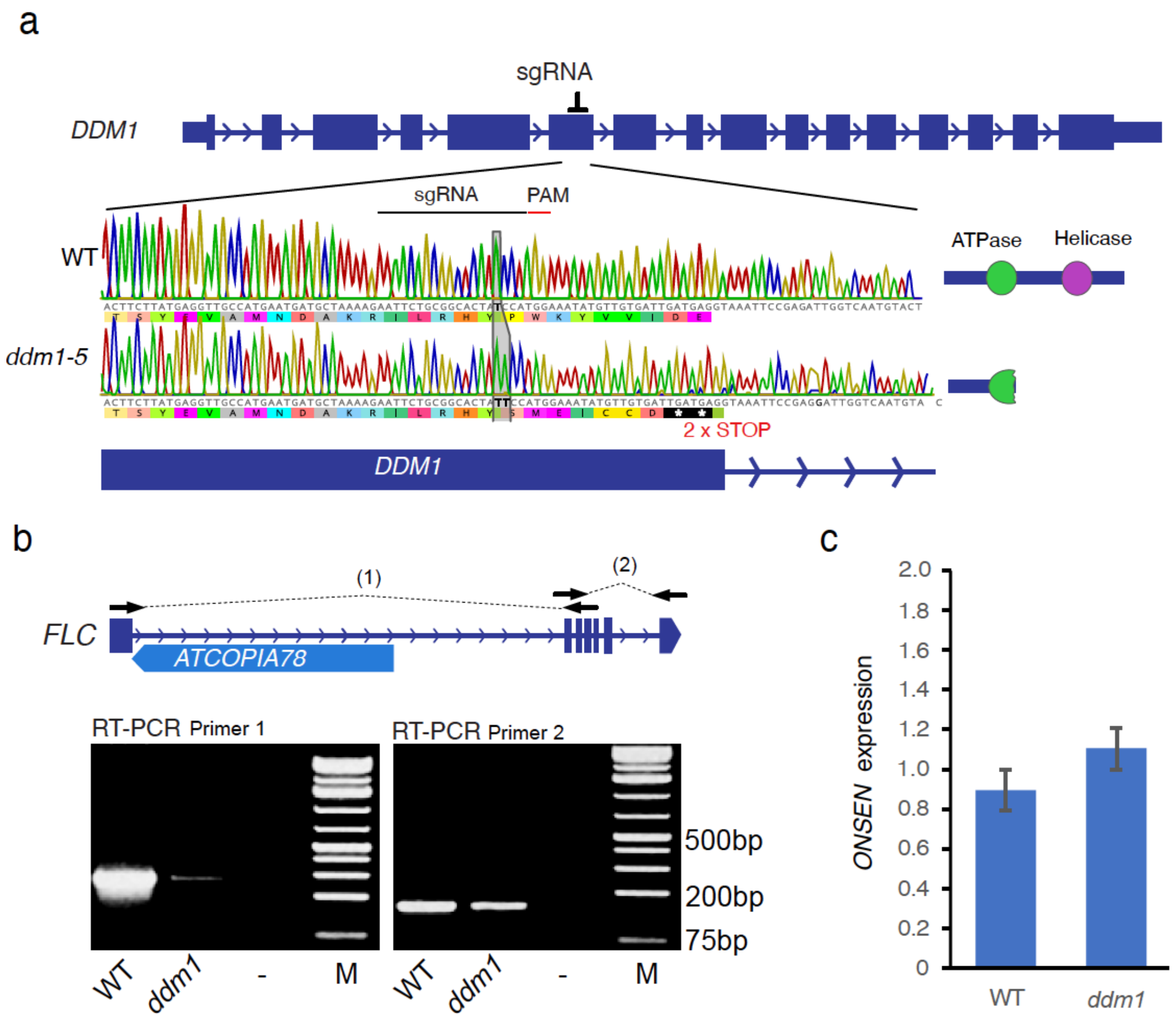


**Supplementary Figure 5. CRISPR-CAS9 Ag-0 *ddm1* plants. a.** Position of sgRNAs and mutation generated in *DDM1* gene by CRIPR-CAS9. **b**. end-point RT-PCR evaluating the splicing efficiency of intron 1 (primers 1) and 3' (primers 2) end of *FLC* transcripts in wild-type (WT) and *ddm1*. Position of primers 1 and 2 are indicated by arrows. **c**. Relative expression levels of ONSEN. Data are mean ± s.d. (n>10 samples, two biological experiments) and statistical significance for differences was obtained using the MWU test.

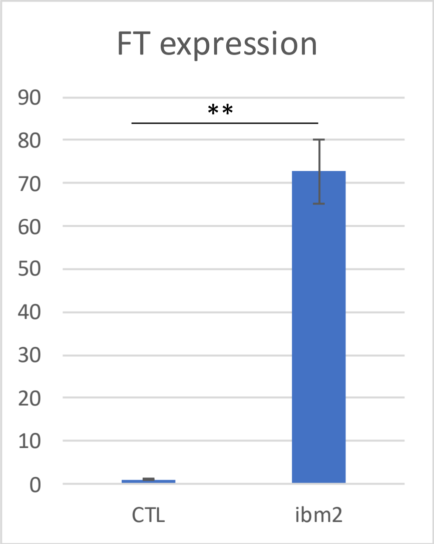


**Supplementary Figure 6. FT expression in Ag-0 ctl. and *IBM2*$^{-/-}$ plants.** Relative expression levels of *FT*. Data are mean ± s.d. (n>10 samples, two biological experiments). ** indicates a statistical significance difference <0.001 obtained using the MWU test.

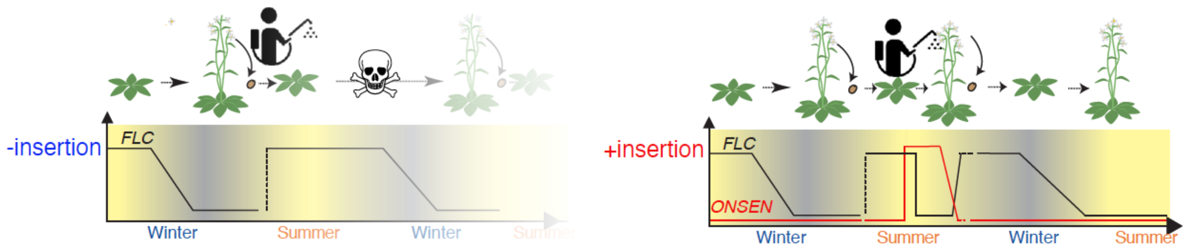


**Supplementary Figure 7. Model summarizing the molecular mechanism underlying the retrotransposon-driven adaptive response to herbicides in Arabidopsis.**

### table S1. Arabidopsis accessions nearby Argentat

| Accession | Latitud | Longitud | nearby city | Collection_year | +Ins | -Ins |
|---|---|---|---|---|---|---|
| Gi-1-1 | 45.0177 | 1.48121 | Gignac Cressensac | 2021 | | X |
| Gi-1-2 | 45.0177 | 1.48121 | Gignac Cressensac | 2021 | | X |
| Gi-1-3 | 45.0177 | 1.48121 | Gignac Cressensac | 2021 | | X |
| Gi-2-2 | 45.0195 | 1.47937 | Gignac Cressensac | 2021 | | X |
| Gi-2-3 | 45.0195 | 1.47937 | Gignac Cressensac | 2021 | | X |
| Arg-4-1 | 44.9965 | 1.30142 | Archignac | 2021 | | X |
| Arg-4-2 | 44.9965 | 1.30142 | Archignac | 2021 | X | |
| Arg-4-4 | 44.9965 | 1.30142 | Archignac | 2021 | X | |
| Gi-2-1 | 45.0195 | 1.47937 | Gignac Cressensac | 2021 | | X |
| Gi-3-3 | 45.0182 | 1.48076 | Gignac Cressensac | 2021 | | X |
| Tur-1-1 | 45.039153 | 1.600121 | Turenne Gare | 2021 | | X |
| Tur-1-10 | 45.039153 | 1.600121 | Turenne Gare | 2021 | | X |
| Tur-1-11 | 45.039153 | 1.600121 | Turenne Gare | 2021 | X | |
| Tur-1-15 | 45.039153 | 1.600121 | Turenne Gare | 2021 | | X |
| Tur-1-2 | 45.039153 | 1.600121 | Turenne Gare | 2021 | X | |
| Tur-1-20 | 45.039153 | 1.600121 | Turenne Gare | 2021 | | X |
| Tur-1-22 | 45.039153 | 1.600121 | Turenne Gare | 2021 | | X |
| Tur-1-3 | 45.039153 | 1.600121 | Turenne Gare | 2021 | | X |
| Tur-1-4 | 45.039153 | 1.600121 | Turenne Gare | 2021 | | X |
| Tur-1-5 | 45.039153 | 1.600121 | Turenne Gare | 2021 | X | |
| Tur-1-7 | 45.039153 | 1.600121 | Turenne Gare | 2021 | X | |
| Tur-2-10 | 45.0407 | 1.59936 | Turenne Gare | 2021 | X | |
| Tur-2-11 | 45.0407 | 1.59936 | Turenne Gare | 2021 | X | |
| Tur-2-12 | 45.0407 | 1.59936 | Turenne Gare | 2021 | | X |
| Tur-2-13 | 45.0407 | 1.59936 | Turenne Gare | 2021 | X | |
| Tur-2-14 | 45.0407 | 1.59936 | Turenne Gare | 2021 | X | |
| Tur-2-16 | 45.0407 | 1.59936 | Turenne Gare | 2021 | X | |
| Tur-2-20 | 45.0407 | 1.59936 | Turenne Gare | 2021 | | X |
| Tur-2-21 | 45.0407 | 1.59936 | Turenne Gare | 2021 | X | |
| Tur-2-3 | 45.0407 | 1.59936 | Turenne Gare | 2021 | | X |
| Tur-2-5 | 45.0407 | 1.59936 | Turenne Gare | 2021 | | X |
| Tur-2-6 | 45.0407 | 1.59936 | Turenne Gare | 2021 | | X |
| Tur-3-2 | 45.0585 | 1.59181 | Turenne | 2021 | X | |
| Tur-3-3 | 45.0585 | 1.59181 | Turenne | 2021 | | X |
| Tur-3-5 | 45.0585 | 1.59181 | Turenne | 2021 | X | |
| Tur-3-6 | 45.0585 | 1.59181 | Turenne | 2021 | | X |
| Tur-3-8 | 45.0585 | 1.59181 | Turenne | 2021 | X | |
| Tur-4-3 | 45.058 | 1.59213 | Turenne | 2021 | X | |
| Tur-5-2 | 45.0588 | 1.59158 | Turenne | 2021 | | X |
| Tur-5-8 | 45.0588 | 1.59158 | Turenne | 2021 | | X |
| Tur-1 | 45.039153 | 1.600121 | Turenne | 2019 | | X |
| Tur-5 | 45.039153 | 1.600121 | Turenne | 2019 | | X |

| | | | | | | |
|---|---|---|---|---|---|---|
| Tur-8 | 45.039900 | 1.599783 | Turenne | 2019 | X | |
| Tur-9 | 45.039900 | 1.599783 | Turenne | 2019 | | X |
| Tur-12 | 45.039900 | 1.599783 | Turenne | 2019 | X | |
| Tur-13 | 45.039900 | 1.599783 | Turenne | 2019 | | X |
| Tur-17 | 45.040274 | 1.599587 | Turenne | 2019 | | X |
| Tur-18 | 45.040274 | 1.599587 | Turenne | 2019 | | X |
| Cha-1 | 45.133473 | 1.521586 | Champ | 2019 | | X |
| Cha-2 | 45.133473 | 1.521586 | Champ | 2019 | | X |
| Cha-3 | 45.133473 | 1.521586 | Champ | 2019 | | X |
| Cha-6 | 45.136698 | 1.525773 | Champ | 2019 | | X |
| Ag-2 | 45.0917 | 1.93325 | Argentat | 2019 | | X |
| Ag-3 | 45.0917 | 1.93325 | Argentat | 2019 | | X |
| Ag-5 | 45.0917 | 1.93325 | Argentat | 2019 | | X |
| Ag-6 | 45.0917 | 1.93325 | Argentat | 2019 | | X |
| Ag-9 | 45.091949 | 1.932473 | Argentat | 2019 | | X |
| Ag-10 | 45.091949 | 1.932473 | Argentat | 2019 | | X |
| Sch-7 | 45.124463 | 1.895981 | Saint-Chamant | 2019 | | X |
| Sch-8 | 45.124463 | 1.895981 | Saint-Chamant | 2019 | | X |
| Sch-13 | 45.124463 | 1.895981 | Saint-Chamant | 2019 | | X |
| Sch-16 | 45.125018 | 1.895108 | Saint-Chamant | 2019 | | X |
| Arg-4 | 45.125018 | 1.895108 | Archignac | 2019 | | X |
| Arg-6 | 45.125018 | 1.895108 | Archignac | 2019 | X | |
| Arg-8 | 45.125018 | 1.895108 | Archignac | 2019 | | X |
| Arg-10 | 45.125018 | 1.895108 | Archignac | 2019 | X | |
| Arg-11 | 45.125018 | 1.895108 | Archignac | 2019 | | X |
| Arg-13 | 45.125018 | 1.895108 | Archignac | 2019 | | X |
| Arg-14 | 45.125018 | 1.895108 | Archignac | 2019 | | X |
| Arg-15 | 45.125018 | 1.895108 | Archignac | 2019 | | X |
| Arg-16 | 45.125018 | 1.895108 | Archignac | 2019 | | X |
| Arg-17 | 45.125018 | 1.895108 | Archignac | 2019 | | X |
| Arg-18 | 45.125018 | 1.895108 | Archignac | 2019 | | X |
| Mde-0 | 45.079402 | 1.612115 | La Rougerie | 2019 | | X |

**table S2. Arabidopsis populations collected across the south of France (Frachon et al. 2018)**

| pop | LAT | LONG | site | +Ins | -Ins |
|---|---|---|---|---|---|
| COMT-A | 44.540652 | 2.602245 | road verge | x | |
| CRAN-A | 44.529845 | 2.260486 | cementery | x | |
| BESS-A | 44.526359 | 2.730092 | backyard | x | |
| JULI-A | 44.5226 | 2.3635 | field margin | x | |
| NAUV-A | 44.520751 | 2.427404 | wild | x | |
| NAUV-B | 44.520418 | 2.427129 | brownfield | | x |
| NAUV-C | 44.520397 | 2.42721 | sidewalk | x | |
| BELC-C | 44.389212 | 2.336636 | sidewalk | x | |
| BELC-A | 44.387532 | 2.336117 | railway | x | |
| BELC-B | 44.387527 | 2.336782 | sidewalk | | x |
| COLO-C | 44.34806 | 2.339698 | sidewalk | x | |
| COLO-B | 44.34773 | 2.339715 | sidewalk | x | |
| COLO-A | 44.346915 | 2.340243 | sidewalk | x | |
| RADE-A | 44.345163 | 2.620821 | backyard | x | |
| BARA-C | 44.270842 | 2.427551 | railway | x | |
| BARA-B | 44.269727 | 2.426322 | railway | x | |
| LAUZ-A | 44.25608 | 1.140526 | parking lot | | x |
| NAZA-A | 44.220329 | 1.064953 | sidewalk | | x |
| BARR-A | 44.202421 | 1.767492 | backyard | x | |
| DECA-A | 44.199896 | 1.77189 | sidewalk | | x |
| RAYR-B | 44.196006 | 2.493076 | brownfield | x | |
| RAYR-A | 44.196005 | 2.493157 | brownfield | x | |
| CASS-A | 44.17653 | 2.518164 | parking lot | x | |
| NAYR-A | 44.161368 | 2.544711 | road verge | | x |
| PAMP-B | 44.124876 | 2.255184 | road verge | x | |
| PAMP-A | 44.124864 | 2.255514 | wild | | x |
| MONE-A | 44.115354 | 2.094725 | road verge | x | |
| CAPE-A | 44.108545 | 1.990168 | field margin | x | |
| MOUL-A | 44.089762 | 2.296094 | railway | x | |
| PANA-C | 44.078884 | 2.711136 | cementery | x | |
| DIEU-A | 44.059797 | 1.220975 | sidewalk | | x |
| VICT-B | 44.052243 | 2.834023 | field | | x |
| VICT-C | 44.052243 | 2.834023 | field | | x |

| | | | | | |
|---|---|---|---|---|---|
| ROME-A | 44.041553 | 2.909576 | parking lot | | x |
| BROU-C | 44.03326 | 2.638683 | field | | x |
| BROU-B | 44.0331 | 2.6387 | field margin | x | |
| VALE-A | 44.022296 | 2.403434 | cementery | x | |
| CERN-B | 44.014684 | 2.967927 | railway | x | |
| CERN-A | 44.01194 | 2.966488 | railway | | x |
| BROU-A | 43.999349 | 2.621684 | brownfield | | x |
| ANGE-B | 43.91214 | 1.656855 | cementery | x | |
| ANGE-A | 43.911999 | 1.656649 | cementery | x | |
| LECT-A | 43.911721 | 0.629745 | railway | x | |
| LECT-B | 43.911721 | 0.629745 | railway | x | |
| GAIL-B | 43.909032 | 1.901077 | railway | x | |
| GAIL-A | 43.908928 | 1.900574 | railway | x | |
| GREZ-A | 43.8769 | 0.3497 | field | | x |
| TARN-C | 43.85328 | 1.502009 | sidewalk | x | |
| MONT-B | 43.852723 | 1.873536 | field margin | x | |
| MONT-A | 43.852212 | 1.87432 | field margin | x | |
| REAL-A | 43.83165 | 2.20155 | road verge | x | |
| CAMA-E | 43.8249 | 2.8817 | cementery | | x |
| CAMA-C | 43.824878 | 2.881661 | cementery | | x |
| CAMA-D | 43.823736 | 2.881003 | cementery | | x |
| PASD-B | 43.811758 | 2.871661 | wild | | x |
| FAYA-A | 43.8021 | 2.9517 | cementery | | x |
| LAGR-A | 43.7953 | 1.0738 | sidewalk | x | |
| BELL-A | 43.790307 | 1.106456 | sidewalk | x | |
| LUZE-B | 43.7644 | 1.7536 | field | | x |
| CEPE-A | 43.755183 | 1.435978 | silo site | x | |
| AMBR-A | 43.733229 | 1.823869 | sidewalk | x | |
| MERV-B | 43.725141 | 1.247629 | road verge | x | |
| MERV-A | 43.720426 | 1.296824 | sidewalk | x | |
| PREI-A | 43.717856 | 0.623298 | railway | x | |
| CAST-A | 43.698534 | 1.427856 | cementery | x | |
| ESPE-B | 43.693335 | 2.534582 | backyard | x | |
| ROQU-B | 43.667907 | 2.290214 | cementery | x | |
| MARS-B | 43.662542 | 0.718265 | silo site | x | |
| MARS-A | 43.6625 | 0.7183 | silo site | x | |

| | | | | | |
|---|---|---|---|---|---|
| FERR-A | 43.657743 | 2.44371 | sidewalk | x | |
| DAMI-A | 43.654515 | 1.977636 | railway | x | |
| DAMI-B | 43.654515 | 1.977636 | railway | x | |
| DAMI-C | 43.654515 | 1.977636 | railway | x | |
| VIEL-A | 43.623801 | 2.089616 | railway | x | |
| MONF-A | 43.616254 | 0.972435 | railway | x | |
| SALV-A | 43.602578 | 2.36338 | road verge | x | |
| LOUB-B | 43.574647 | 1.785723 | sidewalk | x | |
| LOUB-A | 43.574273 | 1.786038 | sidewalk | x | |
| LANT-B | 43.564943 | 1.65239 | silo site | x | |
| LANT-D | 43.564822 | 1.65201 | silo site | x | |
| LANT-C | 43.564822 | 1.65201 | silo site | x | |
| LABR-A | 43.5312 | 2.2626 | sidewalk | x | |
| AUZE-A | 43.527792 | 1.491628 | sidewalk | x | |
| THOM-A | 43.513975 | 1.082859 | sidewalk | | x |
| MAZA-A | 43.497754 | 2.375372 | railway | x | |
| SAMA-A | 43.494325 | 0.92391 | road verge | | x |
| MEDA-A | 43.490485 | 0.461439 | sidewalk | x | |
| LAMA-A | 43.487424 | 1.243559 | field | | x |
| SEIS-A | 43.4873 | 0.588 | sidewalk | x | |
| LAMA-B | 43.4797 | 1.2416 | wild | | x |
| AURE-B | 43.477976 | 1.452214 | cementery | x | |
| SAUB-C | 43.475583 | 1.367589 | backyard | | x |
| SAUB-B | 43.474107 | 1.364175 | backyard | | x |
| MONB-A | 43.46529 | 0.986273 | road verge | | x |
| CLAR-B | 43.465281 | 1.218577 | sidewalk | | x |
| SAUB-A | 43.4649 | 1.3651 | sidewalk | x | |
| CLAR-A | 43.464776 | 1.219019 | field | | x |
| CLAR-C | 43.464058 | 1.21799 | sidewalk | | x |
| VILLE-A | 43.4582 | 1.381 | sidewalk | x | |
| VILLA-A | 43.458174 | 1.380951 | sidewalk | | x |
| LABA-D | 43.458019 | 1.381137 | sidewalk | x | |
| LABA-B | 43.458 | 1.3811 | sidewalk | x | |
| BAZI-A | 43.453602 | 1.620674 | railway | x | |
| SORE-A | 43.452628 | 2.072476 | sidewalk | x | |
| LABA-A | 43.45155 | 1.400498 | sidewalk | | x |

| | | | | | |
|---|---|---|---|---|---|
| LABA-C | 43.451451 | 1.39935 | sidewalk | | x |
| SIMO-A | 43.449392 | 0.734601 | sidewalk | | x |
| JUZE-A | 43.448838 | 1.79053 | sidewalk | x | |
| VILLE-B | 43.440342 | 1.669595 | sidewalk | x | |
| VILLE-C | 43.440024 | 1.670111 | backyard | x | |
| VILLE-D | 43.439733 | 1.670712 | cementery | | x |
| MASS-A | 43.437536 | 0.579271 | road verge | | x |
| MONTI-A | 43.389383 | 0.67282 | backyard | x | |
| MONTI-D | 43.3839336 | 0.67257 | field | | x |
| MONTI-B | 43.3839336 | 0.67257 | field | x | |
| PUYM-B | 43.3729 | 0.7657 | sidewalk | | x |
| BARC-A | 43.362044 | 0.387723 | cementery | | x |
| LUNA-A | 43.339706 | 0.689839 | parking lot | | x |
| BACC-B | 43.3122 | 1.5152 | cementery | | x |
| BACC-C | 43.3122 | 1.5152 | cementery | | x |
| BACC-E | 43.3122 | 1.5152 | cementery | | x |
| BAGNB-A | 43.3122 | 1.5152 | cementery | | x |
| CINT-B | 43.305611 | 1.520735 | parking lot | | x |
| CINT-A | 43.305466 | 1.520441 | railway | x | |
| BOULO-A | 43.28908 | 0.639795 | cementery | x | |
| VILLEM-A | 43.273815 | 0.321238 | field margin | | x |
| MART-A | 43.202147 | 1.010976 | road verge | | x |
| AULO-A | 43.190552 | 0.815774 | sidewalk | | x |
| BERNA-A | 43.16215 | 0.111398 | field margin | | x |
| CARL-A | 43.151102 | 1.3923 | parking lot | x | |
| MONTB-A | 43.130495 | 1.269927 | field margin | x | |
| MONTG-D | 43.127713 | 0.110633 | parking lot | x | |
| MONTG-B | 43.12729 | 0.110681 | wild | x | |
| LESP-A | 43.094237 | 1.719981 | sidewalk | | x |
| BAGNB-B | 43.076454 | 0.151533 | parking lot | | x |
| BANI-B | 43.043644 | 0.234303 | sidewalk | | x |
| BANI-C | 43.043644 | 0.234303 | sidewalk | | x |
| BANI-A | 43.042867 | 0.234732 | sidewalk | | x |
| BULA-A | 43.039803 | 0.277297 | cementery | | x |
| SALE-A | 43.024966 | 0.965965 | backyard | x | |
| CHEI-A | 43.013708 | 0.86707 | field | x | |

| | | | | | |
|---|---|---|---|---|---|
| LABAS-B | 43.008716 | 1.420053 | cementery | x | |
| LACR-C | 43.000155 | 1.075624 | sidewalk | x | |
| LACR-A | 42.999869 | 1.075659 | sidewalk | x | |
| JUZET-A | 42.977713 | 0.756373 | road verge | | x |
| JUZET-B | 42.977354 | 0.755498 | sidewalk | x | |
| JUZET-C | 42.977354 | 0.755498 | sidewalk | x | |
| CASTI-A | 42.920498 | 1.034063 | sidewalk | | x |
| BULE-B | 42.91058 | 1.248122 | sidewalk | | x |
| JACO-C | 42.905839 | 1.406513 | cementery | | x |
| JACO-A | 42.905839 | 1.406513 | cementery | x | |
| SAUR-A | 42.8898 | 1.4852 | field | | x |
| MONTM-A | 42.86156 | 0.595943 | sidewalk | | x |
| MONTM-B | 42.861218 | 0.596869 | sidewalk | x | |
| CIER-C | 42.860166 | 0.601088 | wild | | x |
| CIER-B | 42.859978 | 0.600413 | cementery | | x |
| CIER-A | 42.85332 | 0.602039 | railway | | x |
| CAZA-B | 42.831484 | 0.420091 | road verge | | x |
| LUZE-E | 42.764683 | 1.752959 | railway | | x |
| LUZE-A | 42.764683 | 1.752959 | railway | x | |
| LUZE-D | 42.764419 | 1.753595 | railway | | x |
| AXLE-B | 42.724588 | 1.833497 | parking lot | | x |
| AXLE-A | 42.724197 | 1.834034 | brownfield | | x |
| MERE-A | 42.656618 | 1.836221 | road verge | | x |
| MERE-B | 42.656546 | 1.836175 | road verge | | x |

**table S3. Primers used in this study**

| Name | Sequence 5'-3' |
|---|---|
| FLC-3’-end-spliced F | AGCCAAGAAGACCGAACTCA |
| FLC-3’-end-spliced R | TTTGTCCAGCAGGTGACATC |
| FLC_ex1_ex5_F | CATCCGTCGCTCTTCTCGTC |
| FLC_ex1_ex5_R | TCTAGTCACGGAGAGGGCAG |
| COOLAIR F | TGTATGTGTTCTTCACTTCACTTCTGTCAA |
| COOLAIR R | GCCGTAGGCTTCTTCACTGT |
| FT F | CTTGGCAGGCAAACAGTGTATGCAC |
| FT R | GCCACTCTCCCTCTGACAATTGTAGA |
| SOC1 F | AGCTGCAGAAAACGAGAAGCTCTCTG |
| SOC1 R | GGGCTACTCTCTTCATCACCTCTTCC |
| ATCOPIA 78 F | CCACAAGAGGAACCAACGAA |
| ATCOPIA 78 R | TTCGATCATGGAAGACCGG |
| AGL 24 F | GAGGCTTTGGAGACAGAGTCGGTGA |
| AGL 24 R | AGATGGAAGCCCAAGCTTCAGGAA |
| FLM (beta) F | CATGCTGATGAACTTAGAGCCTTAGATC |
| FLM (beta) R | CAGCAACGTATTCTTTCCCAT |
| PP2A F | TATCGATGACGATTCTTCGTGCAG |
| PP2A R | GCTTGGTCGACTATCGGAATGAGAG |
